# Global gut content data synthesis and phylogeny delineate reef fish trophic guilds

**DOI:** 10.1101/2020.03.04.977116

**Authors:** Valeriano Parravicini, Jordan M. Casey, Nina M. D. Schiettekatte, Simon J. Brandl, Chloé Pozas-Schacre, Jérémy Carlot, Graham J. Edgar, Nicholas A. J. Graham, Mireille Harmelin-Vivien, Michel Kulbicki, Giovanni Strona, Rick D. Stuart-Smith, Jason Vii

**Author notes:** these authors contributed equally (Valeriano Parravicini, Jordan M. Casey).

## Abstract

The diversity of life on our planet has produced a remarkable variety of biological traits that characterize different species. Such traits are widely employed instead of taxonomy to increase our understanding of biodiversity and ecosystem functioning. However, for species’ trophic niches, one of the most critical aspects of organismal ecology, a paucity of empirical information has led to inconsistent definitions of trophic guilds based on expert opinion. Using coral reef fishes as a model, we show that experts often disagree on the assignment of trophic guilds for the same species. Even when broad categories are assigned, 60% of the evaluated trait schemes disagree on the attribution of trophic categories for at least 20% of the species. This disagreement greatly hampers comparability across studies. Here, we introduce a quantitative, unbiased, and fully reproducible framework to define species’ trophic guilds based on empirical data. First, we synthesize data from community-wide visual gut content analysis of tropical coral reef fishes, resulting in trophic information from 13,961 individuals belonging to 615 reef fish species across all ocean basins. We then use network analysis to cluster the resulting global bipartite food web into distinct trophic guilds, resulting in eight trophic guilds, and employ a Bayesian phylogenetic model to predict trophic guilds based on phylogeny and maximum body size. Our model achieved a misclassification error of 5%, indicating that our approach results in a quantitative and reproducible trophic categorization scheme, which can be updated as new information becomes available. Although our case study is for reef fishes, the most diverse vertebrate consumer group, our approach can be applied to other organismal groups to advance reproducibility in trait-based ecology. As such, our work provides an empirical and conceptual advancement for trait-based ecology and a viable approach to monitor ecosystem functioning in our changing world.

## Introduction

A fundamental goal in ecology is to understand the mechanisms behind the maintenance of biodiversity and ecosystem functioning [1,2]. Understanding the ecological niches of species is central to this endeavor [3,4]. In fact, the degree of niche overlap among species can be a major determinant of the positive relationship among species richness [5], ecosystem productivity [6–8], and ecosystem vulnerability [9] since limited functional redundancy can make ecosystems more prone to lose entire energetic pathways [10–12]. With growing threats to biodiversity, the need to quantify the impact of biodiversity loss has amplified the use of functional groups, which are species groups that share common ecological characteristics and are often defined with coarse, categorical descriptors of species traits [13–16].

Delineating the ecological niche with discrete categories has several operational advantages. First, grouping species into categories helps decompose highly complex ecosystems into comprehensible units, while traditional taxonomic analyses may be difficult to interpret in highly diverse ecosystems. Second, ecological predictions tied to individual species are restricted to the geographic range of the species, whereas predictions of functional groups can be globally comparable. Third, the use of functional groups enables the quantification of functional metrics (e.g. functional richness and functional redundancy) from a standard community data matrix without complex experiments [17–19]. The promise of developing “user-friendly” metrics for functional ecology has motivated the employment of trait-based data in community ecology; even with a paucity of empirical information, it is often assumed that experts can achieve accurate descriptions of the ecological niche of species [17,20,21].

Coral reefs, one of the most diverse marine ecosystem on Earth, have inspired a plethora of trait-based ecological studies, with significant recent efforts to compile trait-based datasets for two major components of this ecosystem: corals and fishes [22,23]. For some traits, such as maximum body size in fishes, the compilation process is simple because unidimensional, quantitative data (e.g. maximum total length) are compiled in publicly accessible databases; however, when it comes to species’ diet or behavior, obtaining consensual data is much more difficult. For example, dietary data are multidimensional (i.e. various prey items can be recorded across individuals), ontogenetically variable (i.e. diet differs between juveniles and adults), spatially variable (i.e. species may show dietary plasticity across locations), and prone to methodological differences and observer bias. Therefore, researchers that employ traits to delineate trophic groups or behavioral characteristics commonly rely on expert opinion [19]. While there is some agreement among experts on which traits are relevant (e.g. diet, mobility, body size, diel activity), there is often an implicit disagreement on the necessary categories to describe these traits. For example, across the coral reef literature, the number and resolution of reef fish trophic guilds substantially differs. Studies commonly define three [24] to eight [25] trophic guilds, with particular ambivalence on the resolution at which to define herbivores and invertivores [26–29].

Among all trait classification schemes for reef fishes, only a few are openly accessible. Consequently, different research groups tend to employ proprietary functional classifications, with little possibility to cross-check and compare assigned traits with previous classifications. The classification of species into functional groups has advantages for our understanding of ecological patterns [30,31]. However, the lack of agreement and the limited transparency of trait-based datasets can conjure skepticism and inhibit the emergence of general patterns.

Here, we quantify expert disagreement in the definition of reef fish trophic guilds and propose a novel, transparent, and quantitative framework to delineate trophic guilds. Using coral reef fishes as a case study, we compiled all quantitative, community-wide dietary analyses from several locations across the Pacific and Caribbean and used network analysis to define eight modules that correspond to trophic guilds. We then examined phylogenetic niche conservatism with a phylogenetic Bayesian multinomial model that predicted trophic guilds to the global pool of coral reef fishes, including measures of uncertainty. Our framework is fully reproducible and can be extended and updated as new data become available.

## Materials and Methods

### Assessment of expert agreement

We systematically searched Google Scholar, including papers since 2000, using the following keywords: “coral reefs” AND “reef fish” AND (“fish community” OR “fish assemblage”) AND “diet” AND (“functional group” OR “functional trait” OR “functional entity” OR “trophic guild” OR “trophic group”). The results were individually assessed to find data on trophic guilds. We only considered studies performed at the community level that targeted all trophic levels. Most studies were excluded because they only included specific families or groups, or the data were not provided with the publication. We often found redundant results, with groups publishing several papers using the same classification scheme. In those cases, only the first reference was retained. We contacted authors when trophic classifications were widely used across the literature, but data were not provided with the publications.

Our search yielded a total of eight independent trophic classifications, including Mouillot et al. [26] and Parravicini et al. [27] with 6,316 species, Brandl et al. [32] with 257 species, Halpern and Floeter [33] with 1,046 species, Graham et al. [34] with 126 species, Morais and Bellwood [35] with 515 species, Yeager et al. [36] with 480 species, Newman et al. [37] with 84 species, and Stuart-Smith et al. [29] with 3189 species. The classifications were not uniform in terms of the number and nature of trophic guilds. To achieve comparability, we converted the original classification to match five broad trophic guilds: *herbivores and detritivores*, *invertivores*, *omnivores*, *planktivores*, and *piscivores*. All of the classifications could be reattributed to these categories with the exception of Graham et al. [34], which did not include the category *omnivores*. In this case, the comparison was made only across the four comparable guilds.

In order to assess expert agreement, we compared each possible pair of classifications that shared at least 30 species, generated a confusion matrix, and measured agreement as the proportion of species with matching trophic guild assignments. We then calculated the average agreement between classification pairs for each trophic guild.

### Data collection on fish gut contents

To provide a quantitative definition of trophic guilds for reef fishes, we collected gut content data across the literature at the individual or species level for Chondrichthyes (i.e. cartilaginous fishes) and Osteichthyes (i.e. bony fishes). We obtained dietary information from six published works: Hiatt & Strasburg (1960) for the Marshall Islands [38], Randall (1967) for Puerto Rico and the Virgin Islands [39], Hobson (1974) for Hawaii [40], Harmelin-Vivien (1979) for Madagascar [41], and Sano et al. (1984) for Okinawa [42]. In addition, we provide hitherto unpublished data on the gut contents of 3,015 individuals collected in New Caledonia from 1984 to 2000.

All dietary information was based on visual gut content analysis where prey preference was quantified as volumetric percentage or item frequency. The data were standardized and analyzed as proportions. To our knowledge, the compiled dataset represents the first compilation of detailed coral reef food webs across ocean basins. A total of 13,961 non-empty fish guts belonging to 615 species were analyzed, and more than 1,200 different prey items were described in the original datasets.

First, fish species and family names were taxonomically verified and corrected with the R package *rfishbase* [43]. Only species with at least ten non-empty guts were kept for further analysis. The taxonomic classification of each prey item was then obtained, and all poorly informative (e.g. unidentified fragments, unknown species) and redundant items (e.g. “crustacea fragments” when co-occurring with an item already identified to lower taxonomic level such as “shrimp”) were discarded. Prey identification was highly heterogeneous across the six datasets, differing in taxonomic level and the use of common or scientific names (e.g. crabs *versus* Brachyura). In order to make the six datasets comparable, prey items were grouped into 38 ecologically informative prey groups (Table S1). Items were generally assigned to groups corresponding to their phylum or class. Due to the high diversity and detailed descriptions of crustaceans, they were assigned to the level of order or superorder. Most groups follow official taxonomic classifications except for “detritus,” “inorganic,” and “zooplankton.” In the West Indies dataset [39], items labelled as “Algae & Detritus” were assigned to both of the categories “detritus” and “benthic autotroph,” and the percentage was equally divided in two. The category “zooplankton” includes all eggs and larvae regardless of taxonomy.

### Definition of trophic guilds

After data cleaning, we compiled dietary information for 615 species. Of those species, 516 were present in only one location, 66 were collected in two locations, 25 in three locations, 7 in four locations, and only 1 across five locations. Before running an analysis at the species level, we tested whether there was a strong regional signal for species present across more than one location. We created a quantitative bipartite network where fish species at each location were linked to the 38 prey groups. This network was weighted so that edge weights represent the proportional contribution of each prey group to the diet of a species at a given location.

In order to identify network modules that correspond to reef fish trophic guilds and their preferred prey, we used the maximization of the weighted network modularity based on weighted bipartite networks [44]. Since the modularity maximization algorithm has an initial random step, it may converge to different (although similar) suboptimal solutions each time the analysis is performed, which is common across several optimization algorithms, such as simulated annealing [45]. To guarantee reproducibility and reduce the risk of basing our analysis on an outlier, we performed the modularity maximization 500 times and retained the medoid solution, which was identified as the solution with the highest similarity to the other 499 modules. Similarity was assessed as the variation of information [46]. Overall, 68% of the site × location combinations for the same species belonged to the same module. Therefore, we considered the regional effect to be minor and performed the analysis on the global network, ignoring regional variability and increasing the number of individuals per species.

### Testing for phylogenetic conservatism and predicting trophic guilds

We extracted the phylogenetic position of the 615 species used for the definition of trophic guilds through the Fish Tree of Life [47]. 603 out of 615 species were available in the Fish Tree of Life, but only 535 species had verified phylogenetic information. For the taxa available in the Fish Tree of Life without verified phylogenetic information, we retrieved the pseudo-posterior distribution of 100 synthetic stochastically-resolved phylogenies where missing taxa were placed according to taxonomy using the function *fishtree_complete_phylogeny()* in the R package *fishtree* [48].

We quantified the phylogenetic signal by calculating the phylogenetic statistic δ, which uses a Bayesian approach for discrete variables [49]. The δ statistic can be arbitrarily large with a high level of variation, depending on the number of species and trait levels. To evaluate the significance of the δ statistic, we applied a bootstrapping approach where we quantified δ one hundred times after randomly shuffling the trait values.

We then fitted a multinomial phylogenetic regression to predict fish trophic guild according to phylogeny and body size with the R package *brms* [50]. We used a multinomial logit link function. As such, the probability of a particular trophic guild is computed as follows:

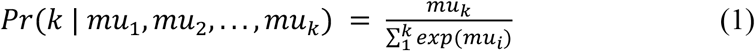

with *muk* defined as:

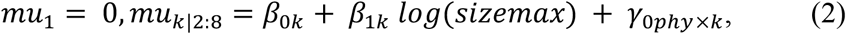

where *β*_0*k*_ is the category-specific fixed-effect intercept, *β*_1*k*_ is the slope for the natural transformed maximum body size for each category *k*, and *γ*_0*phy*×*k*_ is the matrix of random effect coefficients that account for intercept variation based on relatedness as described by the phylogeny for each diet category *k*. We used uninformative priors and ran the model for three chains, each with 6,000 iterations and a warm-up of 1,000 iterations. We visualized the fitted probabilities for each trophic guild with a phylogenetic tree, including the 535 species with verified phylogenetic positions using the R package *ggtree* [49]. Next, we used our model to predict the most likely trophic guild for the global pool of reef fish species. For the extrapolation, we selected all species within reef fish families with more than one representative species (but we also included *Zanclus cornutus*, which is the only species in the family Zanclidae), which resulted in 50 families. Further, we only selected species with a maximum length greater than 3 cm, which was the maximum size of the smallest fish in our compiled database. This selection process resulted in a list of 4,554 reef fish species.

Currently, no streamlined method exists to predict traits for new species from a phylogenetic regression model. We circumvented this issue by extracting draws of the phylogenetic effect (*γ*_0*phy*×*k*_) for each species included in the model. We subsequently predicted the phylogenetic effects for missing species with the help of the function *phyEstimate* from the R package *picante* [51]. This function uses phylogenetic ancestral state estimation to infer trait values for new species on a phylogenetic tree by re-rooting the tree to the parent edge to predict the node. We repeated this inference across 2,000 draws. Per draw, we randomly sampled one of the one hundred trees. Then, we predicted the probability of each species to be assigned to each diet category by combining the predicted phylogenetic effects with the global intercept and slopes for maximum body size for each draw. Finally, we summarized all diet category probabilities per species by taking the mean and standard deviation across all 2,000 draws.

We quantified the total standard deviation (i.e. the square root of the quadratic sum of the standard deviations in each category) and the negentropy value, a measure of certainty calculated by subtracting one from the entropy value (i.e. uncertainty). Thus, the negentropy value lies between 0 and 1, and the higher the value, the higher the certainty for trophic guild assignment (i.e. if a given species has a high probability of assignment to a dietary category, the negentropy value will be high).

## Results

### Assessment of expert agreement

We evaluated the agreement among eight distinct and independent trophic guild classifications by comparing the classification schemes in pairs. Considering the broadness of the expert-assigned categories, we found remarkably low agreement. The median agreement between pairs, expressed as the proportion of species with matching trophic group assignments, was 77% (Fig. 1). For 50% of the pairwise comparisons, at least a quarter of the species were attributed to different trophic groups. In the most severe disagreement, the proportion of mismatched assignments reached 39%. In addition, expert agreement differed depending on the trophic group. Despite a few peaks of disagreement for *herbivores and detritivores* (~20%), overall, there was high agreement among experts for this trophic guild, with an average agreement of 94% (Fig. 1b). On the contrary, *omnivores* showed the highest mismatch, with experts disagreeing on an average of 30% of the species and peaks of disagreement higher than 60% (Fig. 1b).

**Figure 1.**
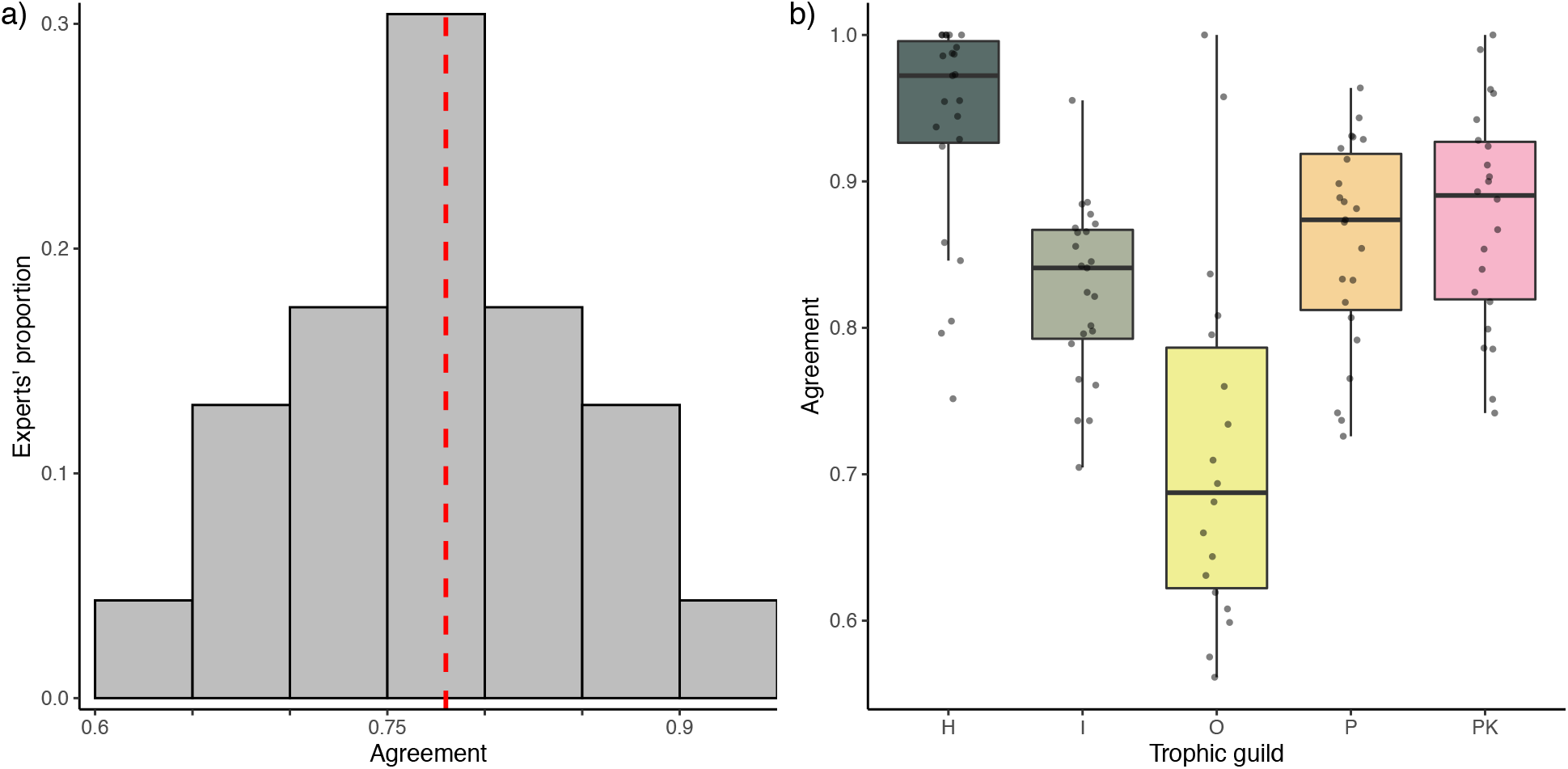
Expert agreement on trophic guild assignment. (a) The distribution of the agreement (i.e. proportion of species assigned to the same trophic category) across the 32 comparisons between pairs of experts. The red dotted line represents the median. (b) Agreement between pairs of experts by trophic category (H=herbivores and detritivores, I=invertivores, O=omnivores, P=piscivores, PK=planktivores).

Expert agreement was variable and often homogeneously distributed around the mean for all the trophic categories. Therefore, the high agreement between a few combinations of experts did not necessarily exclude peaks of disagreement (Fig. 1b). The analysis of individual confusion matrices between pairs of experts revealed high heterogeneity (Fig. 2). For example, Morais and Bellwood [35] were generally in agreement with Mouillot et al. [26] (across 89% of the 515 species in common), while Mouillot et al. [26] agreed with Stuart-Smith et al. [29] across only 68% of the 2211 species in common.

**Figure 2.**
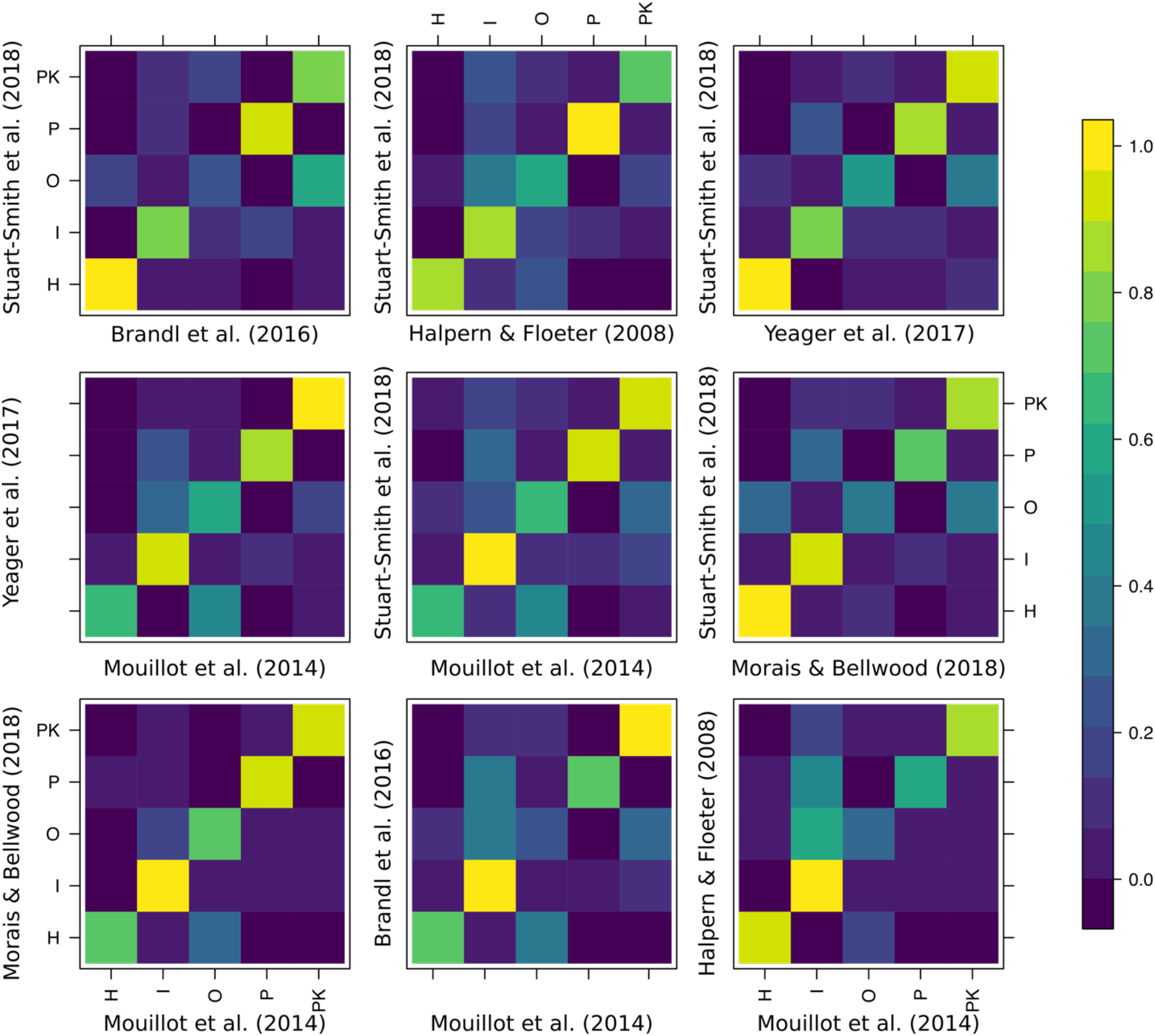
Confusion matrices of the agreement between pairs of experts that share at least 200 species in common. Colors represent proportions of species in each trophic guild as classified by experts (H=herbivores and detritivores, I=invertivores, O=omnivores, P=piscivores, PK=planktivores).

Surprisingly, there was also a high heterogeneity in groups with high disagreement (i.e. multiple alternative assignments for species not assigned to the same trophic group). Species classified as *invertivores* according to one expert were considered *omnivores*, *piscivores*, or *planktivores* according to other classification schemes (Fig. 2). Similarly, species considered *omnivores* by one expert were alternatively considered *invertivores*, *herbivores and detritivores*, or *planktivores* by another expert.

### Definition of trophic guilds

We defined trophic guilds by identifying modules (i.e. combinations of predators and prey) that maximize the weighted modularity of the global network. Our analysis robustly identified eight distinct modules that correspond to different trophic guilds (Fig. 3). We identified these trophic guilds as:

1. *Sessile invertivores*: species predominantly feeding on Asteroidea, Bryozoa, Cirripedia, Porifera, and Tunicata;
2. *Herbivores, microvores*, *and detritivores (HMD)*: species primarily feeding on autotrophs, detritus, inorganic material, foraminifera, and phytoplankton;
3. *Corallivores*: species primarily feeding on Anthozoa and Hydrozoa;
4. *Piscivores*: species primarily feeding on Actinopterygii and Cephalopoda;
5. *Microinvertivores*: species primarily feeding on Annelida, Arachnida, Hemichordata, Nematoda, Peracarida, and Nemertea;
6. *Macroinvertivores*: species primarily feeding on Mollusca and Echinodermata;
7. *Crustacivores*: species primarily feeding on Decapoda and Stomatopoda;
8. *Planktivores*: species mainly feeding on zooplankton and Harpacticoida.

**Figure 3.**
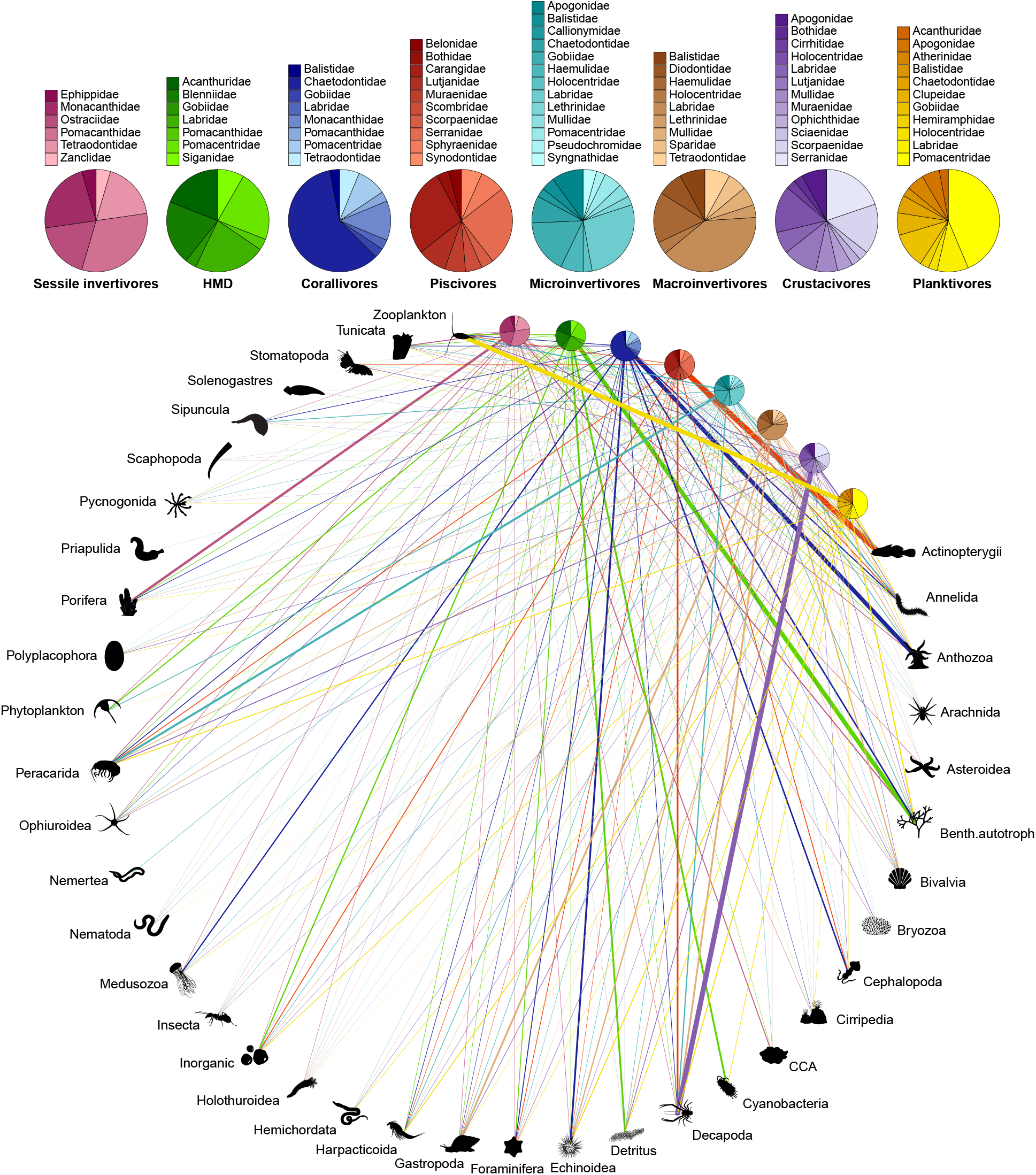
Bipartite network including 615 fish species (grouped into eight trophic guilds; HMD = *herbivores, microvores, and detritivores*) and their prey items (grouped into 38 categories; see Table S1). The relative proportion of each prey category consumed by each trophic guild corresponds with the width of each interaction bar. The pie charts show the relative proportion of fish families within each trophic guild.

### Phylogenetic signal

To evaluate the significance of the phylogenetic statistic value (δ = 9.37), we applied a bootstrapping approach and quantified δ after randomly shuffling the trait values 100 times. The median δ of these null models was 0.000199 (95% confidence interval [0.000196, 0.000204]), indicating a strong phylogenetic signal associated with the eight trophic guilds.

Phylogeny and maximum body size were sufficient to correctly predict the trophic guild of 97% of the species in our dataset. For most families, there was strong phylogenetic conservatism, which resulted in the high confidence of these predictions (Fig. 4). Within some families, however, closely related species displayed distinct dietary preferences. The uncertainty around these family-level predictions was higher, as showcased by high negentropy values for families such as Balistidae, Diodontidae, and Labridae.

**Figure 4.**
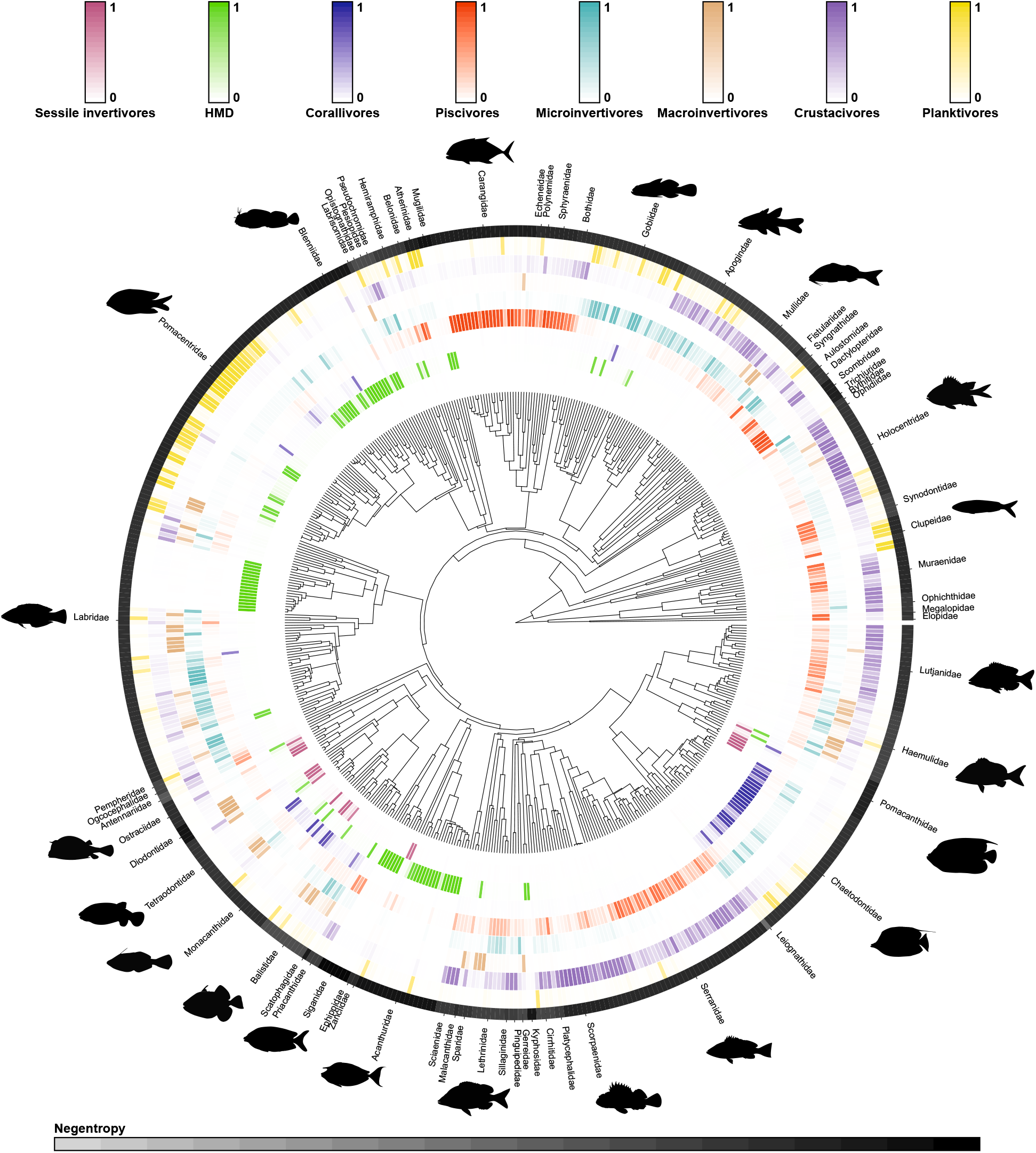
Phylogenetic tree of 535 reef fish species with fitted trophic guild assignments based on empirical dietary data. Trophic guild predictions were made with a Bayesian multinomial phylogenetic regression. The probability of trophic guild assignments for each species is visualized with color scales (depicted above the phylogenetic tree), with darker colors indicating a higher probability of assignment. In the outer black ring, each distinct segment represents a fish family (with silhouettes included for the most speciose families). Uncertainty of overarching trophic guild assignment for each fish family is visualized with negentropy values (i.e. reverse entropy); thus, darker shades indicate a higher degree of certainty of trophic guild assignment.

Given the high predictive performance of our Bayesian phylogenetic model, we used our model to extrapolate the probability of all reef fish species belonging to the eight trophic guilds and assigned the trophic guild with the highest probability. Using leave-one-out cross validation, the final accuracy of this approach was 65%, which is comparable to other phylogenetically-extrapolated traits applications, such as those involving microbial traits [52].

By inspecting the confusion matrix of the leave-one-out cross validation, we obtained more detailed information on the accuracy of the trophic guild predictions (Fig. S1). Most categories were well predicted with our extrapolation approach. In particular, the *sessile invertivores*, *HMD*, and *piscivores* trophic guilds were predicted with high accuracy (77%, 75%, and 73% correct predictions, respectively). The confusion matrix also provided information on incorrectly assigned categories. For example, when *piscivores* were incorrectly assigned, they were mostly classified as *crustacivores*. However, the network plot revealed that the fishes classified as *piscivores* also fed on crustaceans (mostly decapods), so this “incorrect assignment” was grounded in ecological reality and reflected uncertainty within the model. Additionally, the *microinvertivores* trophic guild had the highest proportion of inaccurate predictions (52% correct predictions). Here, species were often misclassified as *crustacivores* or *planktivores*.

## Discussion

To harness the full strength of trait-based approaches, functional ecology requires standardized and reproducible trait classification schemes across taxonomic groups [53–55]. Rather than rely on expert opinion for the assignment of trophic groups, which often results in variable assignments, we demonstrate that the categorization of trophic guilds can be achieved with a quantitative and reproducible framework that is grounded in empirical data across biogeographic regions. We employed network analysis to partition 615 tropical coral reef fish species into eight trophic guilds based on a synthesis of globally distributed, community-wide fish dietary analyses. We then applied a Bayesian phylogenetic model that predicts trophic guilds based on phylogeny and body size and attained a 5% misclassification error. Unlike traditional trophic guilds based on expert opinion [26,27,29,32–37], our trophic classifications are reproducible, provide uncertainty estimates, and can be updated and improved in the future with additional dietary information. By making our trophic classification framework publicly available, we aim to encourage a new, accessible benchmark for the definition of trophic guilds. Given the growing number of trait-based studies that assign trophic guilds to understand and monitor ecosystem functioning in our changing world, it is imperative that we establish comparable and reproducible trophic classification frameworks.

Our findings highlight the discordance of expert opinion in the assignment of trophic guilds and the necessity to develop a quantifiable, reproducible classification scheme that is accessible to the wider scientific community (c.f. [56]). To address this issue, the framework proposed herein represents the first implementation of a quantifiable classification scheme for coral reef fishes, including measures of uncertainty around trophic guild assignments and providing a new path forward to standardize the definition of traits. Despite broad similarities between the trophic guilds reported in the literature and the groups identified by our analysis, our classification scheme reveals a higher level of partitioning among invertebrate-feeding fishes as compared to previously proposed trophic guilds. In the past, invertebrate-feeding fishes were generally considered *sessile invertivores*, *mobile invertivores*, or *omnivores* (e.g. [27,28,36]), but we identify five distinct invertebrate-feeding groups: *corallivores*, *sessile invertivores*, *microinvertivores*, *macroinvertivores*, and *crustacivores*. Given the extreme numerical dominance of invertebrates in coral reef environments [57], the collapse of all invertebrate-feeders into two or three trophic groups was possibly an artefact of expert oversight, and the expansion of invertebrate-feeding trophic guilds to five groups stands to improve ecological resolution of fishes feeding on invertebrate prey.

In contrast to the high resolution achieved within invertebrate-feeding groups, our classification achieved limited resolution among the nominally herbivorous species, *herbivores, microvores, and detritivores* (*HMD*). Across the literature, past classification schemes often separate macroalgal feeders, turf algae croppers, and detritivores (e.g. [26,27]). The lack of precision in our framework is rooted in the difficulty in distinguishing algae, microbes, and detritus within the alimentary tract of fishes, resulting in the pooling of these ingested items during the visual assessment of fish gut contents. Consequently, species classified as *HMD* may have fundamentally different foraging strategies, dietary preferences, and evolutionary histories [58], which can greatly impact their functional role on coral reefs (e.g. [59]). Thus, while our identified trophic guilds promise increased resolution for fishes that consume animal prey, our identified groupings may not adequately capture consumer-producer dynamics on coral reefs. Emerging techniques, such as gut content metabarcoding, may provide the additional resolution needed to further discriminate prey items in this group [60,61]. Alternatively, coupling diet categorization with other traits, such as feeding behavior, may help to pinpoint the variety of feeding modes that exist within the *HMD* trophic guild.

Our results also highlight the necessity of integrating evolutionary history (i.e. phylogenetics) in trait-based ecology (c.f. [62]). Recently, taxonomy and body size have been revealed as important predictors of fish diet composition and size structure [63,64], and in the highest resolution analyses of coral reef fish diet, taxonomic family was a better predictor of fish diet than broad trophic guilds [60]. Given the exceedingly low rate of misclassification error in our predictions of fish trophic guilds, we posit that phylogeny is a critical variable that should be consistently considered in the assignment of trophic guilds. Across a plethora of organismal groups (e.g. birds [65], reptiles [66], fishes [67,68], insects [69], parasites [70], plants [71], etc), phylogenetic niche conservatism has been alternately supported and dismissed. In our case, when examining fish trophic guilds using 38 prey categories, phylogenetic conservatism is readily apparent at the family level, and therefore, it may allow us to extrapolate trophic assignments to closely related consumer species. However, with increasing dietary resolution beyond what is detailed in the present study, phylogenetic signals may weaken [72] since even closely related species may exhibit dietary specialization [60,73]. In the future, with the availability of higher resolution of dietary information, phylogenetic niche conservatism can be easily examined within our framework.

With ongoing environmental and ecological change, a firm grasp on shifts in ecosystem functioning will depend on the reliable assignment of organismal traits [15] and the comparability of trait-based approaches across space, time, and independent studies [54]. Especially in complex, hyperdiverse environments such as coral reefs, it is imperative to standardize how we measure and report these traits to prevent idiosyncratic results based on subjective trait assignments [19,74]. Trophic guilds are among the most commonly applied traits to assess ecosystem functioning because they directly relate to energy and nutrient fluxes across trophic levels. Thus, our standardized framework to quantitatively assign trophic guilds across coral reef fishes represents a major step forward for coral reef functional ecology, while heeding the call for openly-accessible, reproducible trait databases [22,55,75]. As trait-based ecology continues to be used to examine disturbances and implement management strategies, our cohesive and accessible framework to predict reef fish trophic guilds can provide key insights into the trajectory of coral reef communities. Coupling our trophic guild assignment framework with predictive models could spur the emergence of an early detection system to forecast shifts in ecosystem functioning [30].

Further, our results can serve as the foundation for an online platform that permits researchers to collate, update, and utilize trait-based data on coral reef fishes. Similar to current initiatives across the entire tree of life [55], we envision the creation of an online, user-maintained dietary database to facilitate collaboration and traceability in trait-based reef fish research. One challenge will lie in merging visual fish gut content analysis databases with molecular data, such as gut content DNA metabarcoding (e.g. [60]), and biochemical data, such as stable isotope analysis (e.g. [76]), and short-chain fatty acid profiles (e.g. [77]), which indicate nutritional assimilation rather than the simple ingestion of prey items [58]. Despite this challenge, accessibility to a large breadth of reef fish dietary information would improve our framework. Our proposed trophic guilds are model predictions, so they are only as reliable as the underlying dietary data. In addition, these predictions may suffer from extrapolation biases; for example, if limited dietary information exists across species within a taxonomic group, extrapolations to closely related species are more likely to be assigned erroneous trophic guilds. Consequently, an ongoing, extensive compilation of dietary traits across coral reef fishes will continuously improve our predicted trophic guild assignments.

Finally, our proposed framework is not limited to coral reef fishes; indeed, trophic guild assignments can be quantifiable, reproducible, and transparent, with the inclusion of uncertainty metrics, across many organismal groups. However, the standardization of trophic guilds is sorely lacking across the ecological literature [56], especially based on quantitative data (e.g. [78]). We posit that a similar approach can be readily applied across a multitude of organisms and environments. In fact, given the paucity of dietary information available for coral reef fishes in comparison to other organisms, particularly birds and mammals, building rigorous, global trophic classification schemes for many other organisms should be readily achievable within our framework. With a quantitative, transparent trophic classification scheme that can be augmented over time and is applicable across ecological systems, our framework represents a significant advancement for trait-based ecology and a viable approach to monitor ecosystem dynamics into the future [55].

## Acknowledgements

This work was supported by the Centre de Recherche Insulaires et Observatoire de l’Environnement (CRIOBE) in Perpignan, France. We thank all the researchers who have made their fish gut content data and/or trait-based fish trophic guild assignments publicly available for use by the wider scientific community. All the relevant data used to perform the analyses and the results of our extrapolation will be made available once the manuscript is published in a scientific journal.

## Author Contributions

### Conceptualization

Valeriano Parravicini, Jordan M. Casey, Nina M. D. Schiettekatte

### Data curation

Valeriano Parravicini, Jordan M. Casey, Chloé Pozas-Schacre

### Formal analysis

Valeriano Parravicini, Nina M. D. Schiettekatte

### Funding acquisition

Valeriano Parravicini

### Investigation

Valeriano Parravicini, Jordan M. Casey, Nina M. D. Schiettekatte, Simon Brandl, Chloé Pozas-Schacre, Jérémy Carlot, Graham J. Edgar, Nicholas A. J. Graham, Mireille Harmelin-Vivien, Michel Kulbicki, Giovanni Strona, Rick D. Stuart-Smith, Jason Vii

### Methodology

Valeriano Parravicini, Jordan M. Casey, Nina M. D. Schiettekatte, Simon J. Brandl, Chloé Pozas-Schacre

### Supervision

Valeriano Parravicini, Jordan M. Casey

### Visualization

Valeriano Parravicini, Jordan M. Casey, Nina M. D. Schiettekatte, Simon J. Brandl

### Writing - original draft

Valeriano Parravicini, Jordan M. Casey

### Writing - review & editing

Valeriano Parravicini, Jordan M. Casey, Nina M. D. Schiettekatte, Simon J. Brandl, Chloé Pozas-Schacre, Jérémy Carlot, Graham J. Edgar, Nicholas A. J. Graham, Mireille Harmelin-Vivien, Michel Kulbicki, Giovanni Strona, Rick D. Stuart-Smith, and Jason Vii

## Supporting Information

**Table S1.**
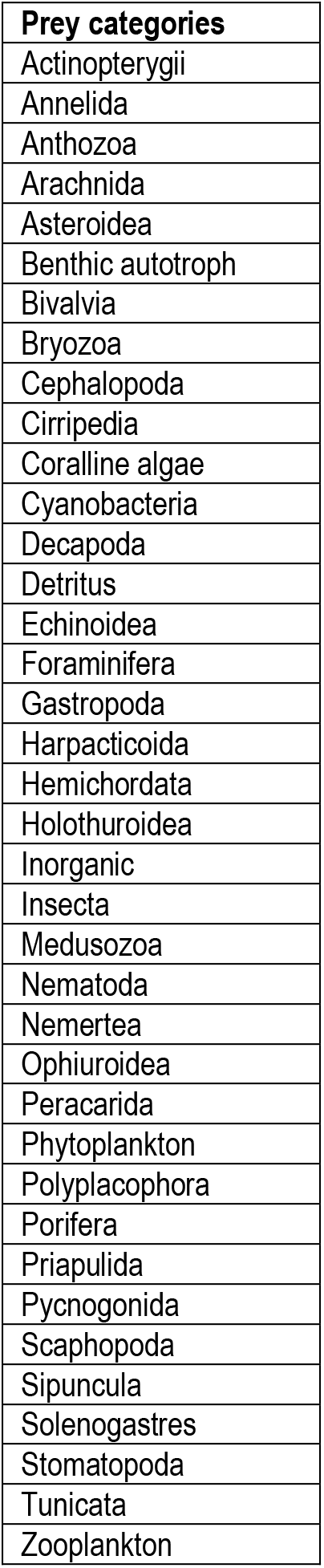
List of prey categories used to define the trophic preferences of coral reef fishes.

**Figure S1.**
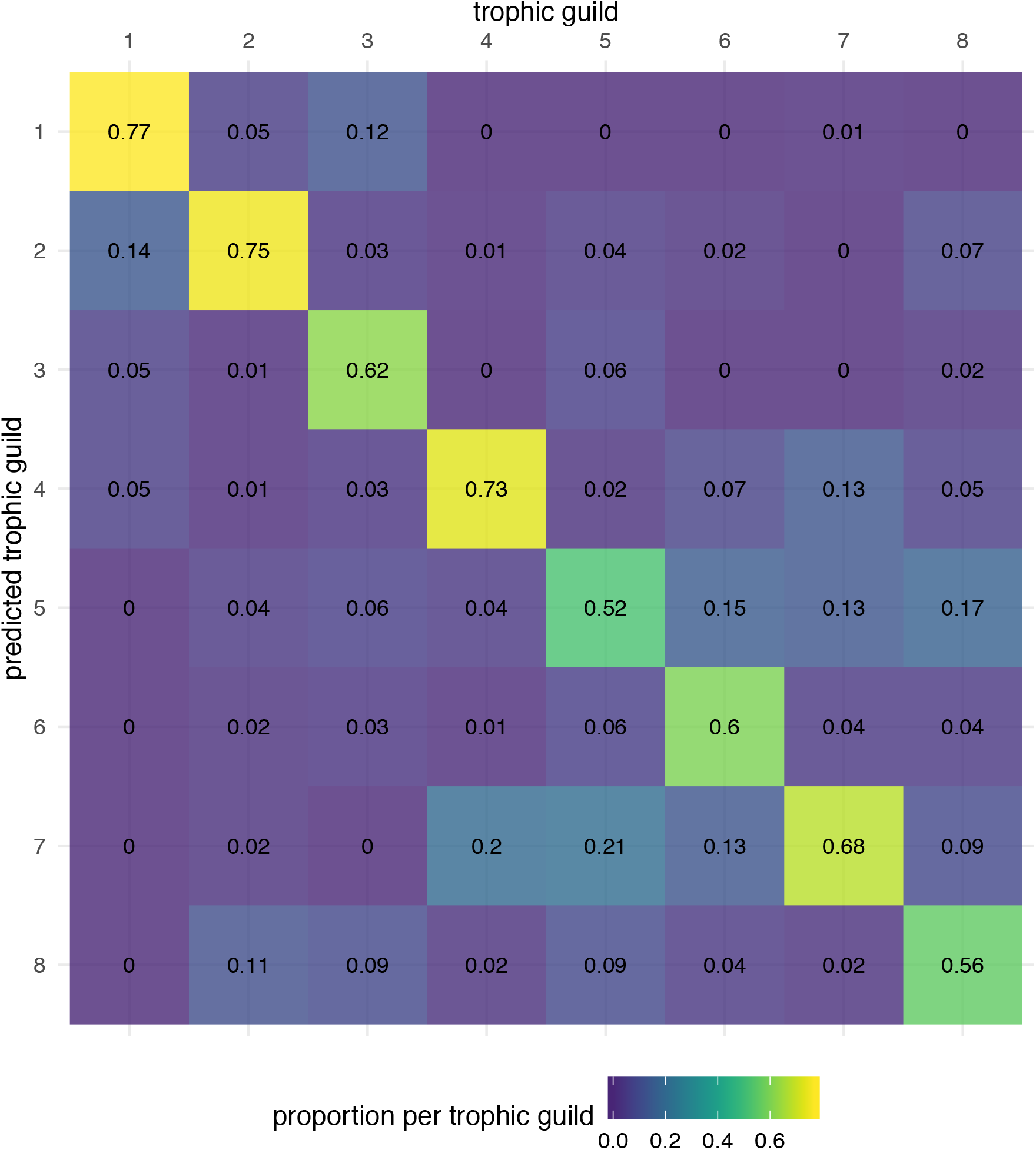
Confusion matrix showcasing the accuracy of the eight trophic guild predictions from the leave-one-out cross validation based on the extrapolation of the Bayesian phylogenetic model. Trophic guilds include: (1) *sessile invertivores*, (2) *herbivores, microvores, and detritivores*, (3) *corallivores*, (4) *piscivores*, (5) *microinvertivores*, (6) *macroinvertivores*, (7) *crustacivores*, and (8) *planktivores*.

